# Best BLAST hit alone cannot be used as evidence of fraud

**DOI:** 10.1101/2021.11.16.468182

**Authors:** Natalia Díaz-Arce, Naiara Rodríguez-Ezpeleta

**Author notes:** Corresponding authors: Natalia Díaz-Arce;, Naiara Rodriguez-Ezpeleta. **ARISING FROM:** Blanco-Fernandez *et al*.: Scientific Reports https://doi.org/10.1038/s41598-021-91020-w (2021).

## Abstract

In a recent study, Blanco-Fernandez, et al. ^1^ applied molecular tools to authenticate fish products and conclude evidence of “worrying international fraud”. They revealed mislabeling in recognizable and unrecognizable fish products labeled as anchovy, hake and tuna commercialized by European companies. Their analyses consisted on extracting DNA from the fish product to be authenticated followed by amplification and sequencing of a suite of DNA markers and on comparing the resulting sequences to the GenBank sequence database using BLAST (Basic Tool Alignment Search Tool) (https://blast.ncbi.nlm.nih.gov/Blast.cgi). By carefully reanalyzing their data, we identify errors in their species identification and conclude that best BLAST hit alone cannot be used as evidence of fraud.

## Introduction

Seafood product traceability is essential to detect intentional or unintentional mislabeling and thus helps reducing unreported, unregulated and illegal fishing while enhancing consumer safety ^2^. Genetic methods have shown to be powerful for seafood product traceability ^3^, in particular, when morphological characteristics cannot be confidently used such as in young age specimens (*e.g*., juveniles of bigeye and yellowfin tunas are very difficult to distinguish ^4^) or in closely related species (*e.g*. black and white anglerfish ^5^), and specially in processed products (*e.g*. filleted and canned), where anatomical traits important for fish identification (*e.g.* head, fins, skin) are absent. Accurate genetic based seafood product traceability requires developing approaches that unequivocally discriminate between species, for which it is essential to understand intra-specific variability as well as each species’ evolutionary context. In their study, Blanco-Fernandez, et al. ^1^ use the best hit of a BLAST search against GenBank to assign species to the sample to be authenticated. Here, by analyzing their BLAST results considering other information, we show that using BLAST alone can lead to erroneous species assignments and thus to conclude fraud when there is not.

## Results and Discussion

One of the striking observations from Blanco-Fernandez *et al*. is the substitution of albacore (*Thunnus alalunga*) by Atlantic bluefin tuna (*Thunnus thynnus*), which is surprising being albacore less than half the price of bluefin tuna (https://www.eumofa.eu/es/home). The authors explain this potential substitution by over-quota-caught bluefin tuna being sold as another species. This is a very strong claim that needs clear evidence to be made. We thus examined the sequences corresponding to those albacore-labelled tuna products (MW557512, MW557513, MW557514) claimed to be mislabelled bluefin tuna due to a best BLAST hit with sequence EU562888, belonging to *T. thynnus* according to GenBank. Our hypothesis was that the mislabeling, rather than in the seafood products, is in the sequence in GenBank due to the mitochondrial introgression reported between *T. alalunga* and *T. thynnus* ^6^. Indeed, it has been estimated that approximately 2-3% of *T. thynnus* individuals have the so-called “alalunga-like” mitochondrial DNA ^7^, which has often misled mitochondrial based phylogenetic inferences of the genus *Thunnus* ^8^. This hypothesis was confirmed by a phylogenetic inference including the putatively mislabelled sequences from Blanco-Fernandez, et al. ^1^. and their best BLAST hit, as well as representative sequences from *T. alalunga*, *T. thynnus* (including those labelled as “alalunga-like”) and *T. albacares* (Figure 1). The tree shows two clearly differentiated clades: one is exclusively composed by *T. thynnus* sequences, while the other includes *T. alalunga*, *T. thynnus* “alalunga-like” sequences as well as the sequences from the putatively mislabelled products and their best BLAST hit. These results refute the mislabeling of *T. thynnus* products as *T. alalunga* reported by Blanco-Fernandez, et al. ^1^. and, more importantly, invalidate their impactful interpretations and conclusions derived therein.

**Figure 1.**
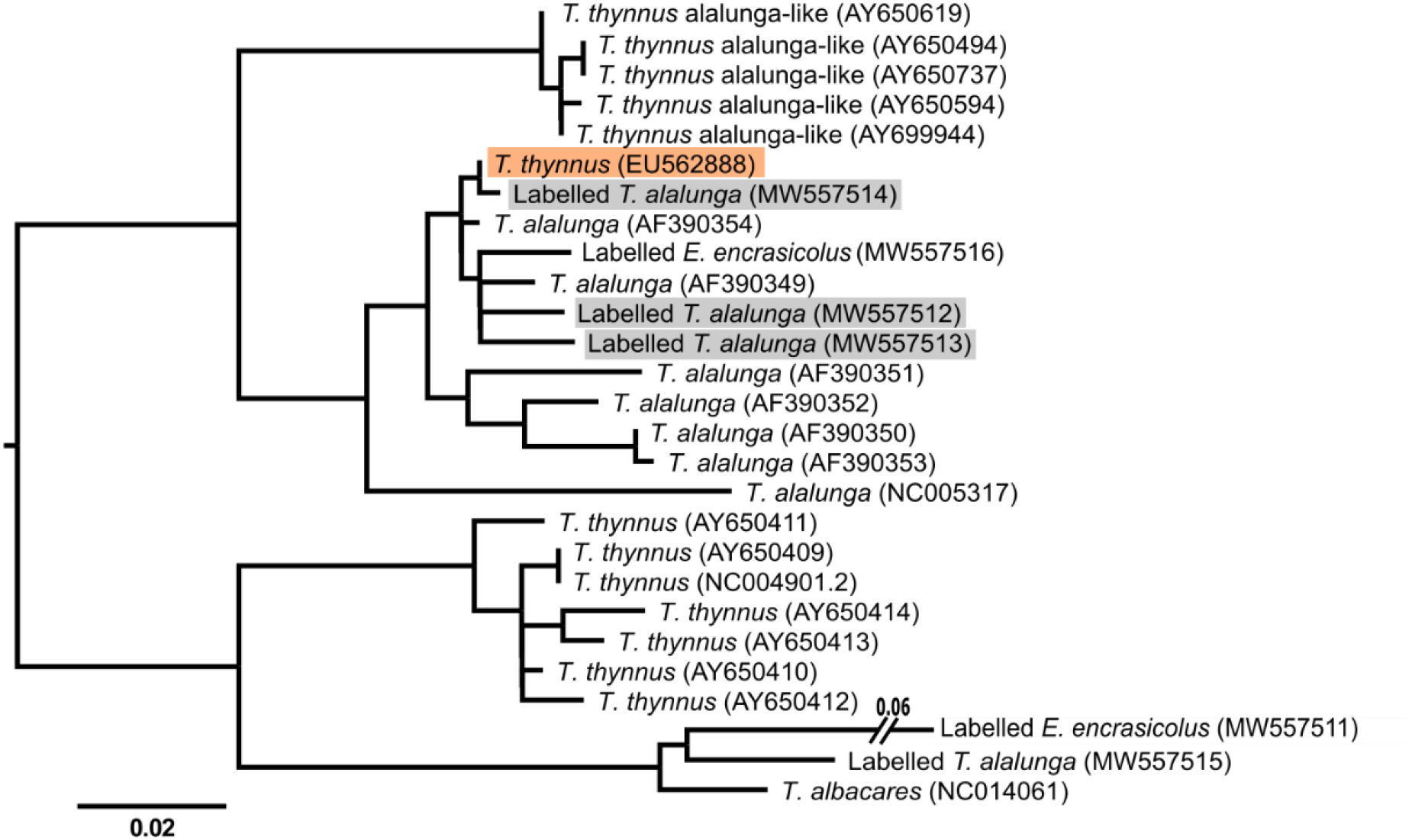
Maximum likelihood phylogenetic inference based on the mitochondrial control region (positions with less than 50% missing data between 5 to 414 of NC_005317 sequence) including all the available tuna identified sequences from Blanco-Fernandez *et al.* (MW557511-16) with the ones labelled *T. alaunga* matching to *T. thynnus* labelled in grey, the best BLAST hit *T. thynnus* sequence, shaded in orange, and reference sequences from *T. albacares*, *T. alalunga*, *T. thynnus* and *T. thynnus* “alalunga-like”). The tree is rooted according to Díaz-Arce, et al. ^8^.

This clear and easy to detect case could be the tip of the iceberg of errors made in food fraud studies relying solely on best BLAST hits to report mislabeling. Indeed, the bluefin tuna and albacore case is not an isolated one, and instances of genetic introgression that could lead to misidentification are increasingly reported in teleost fishes (*e.g*., ^9,10^). Additionally, other factors such as traditionally used morphological characters for species assignment not being diagnostic can also occur and lead to false conclusions regarding mislabelling. This is the case of the black and white anglerfish for which mislabelling was reported ^11^ based on the colour of their peritoneum as species diagnostic character, whereas it has recently been discovered that black anglerfish can have white peritoneum ^12^ and, thus, reported mislabelling was most likely not so.

As shown above, questioning our understanding of the evolutionary history of the species under investigation is essential for seafood fraud studies and could avoid errors such as the one made by Blanco-Fernandez *et al*. when they report mislabeling of albacore products. Additionally, questioning the ability of BLAST searches to distinguish between closely related species could also avoid potential mistakes. Indeed, BLAST results deeply depend on the completeness and accuracy of the reference databases. For example, Blanco-Fernandez, et al. ^1^. reported mislabeling between *Merluccius polli* and *M. paradoxus/capensis* based on best BLAST hit. Yet, this result was obtained thanks to *M. polli* sequences covering their target region being present in the GenBank. Interestingly, had these sequences, which were submitted to GenBank by the same team ^13^, not been available, these *M. paradoxus/capensis* labelled products would have been assigned to *M. merluccius* based on their best BLAST hit. This is due to the next best BLAST hit with other *M. polli* sequences not covering their whole target region (Table 1). Instead, building a phylogenetic tree would have led to the right conclusion in both cases (Figure 2), further highlighting the power of phylogenies over BLAST searches for species assignment.

**Table 1.** Best BLAST hits of sequence MW557504 labelled *M. paradoxus/capensis* ranked according to obtained E value and percentage of identity. Green and yellow indicate best overall BLAST hit if the Machado-Schiaffino, et al. ^13^ sequences (in bold) were not present in GenBank and best *M. polli* BLAST hit. Searches using sequence MW557505 also labeled as *M. paradoxus/capensis* provided virtually identical results.

| Species | Accession | E value | Query Cover | Percent identity |
| --- | --- | --- | --- | --- |
| *Merluccius polli* | **EF362877.1** | 0 | 100% | 100 |
| *Merluccius polli* | **EF362875.1** | 0 | 100% | 99.76 |
| *Merluccius polli* | **EF362876.1** | 0 | 100% | 99.52 |
| *Merluccius paradoxus* | **EF362879.1** | 0 | 100% | 98.32 |
| *Merluccius paradoxus* | **EF362878.1** | 0 | 100% | 98.08 |
| *Merluccius capensis* | **EF362863.1** | 0 | 100% | 97.36 |
| *Merluccius merluccius* | **EF362872.1** | 0 | 100% | 97.12 |
| *Merluccius capensis* | **EF362865.1** | 0 | 100% | 97.12 |
| *Merluccius merluccius* | AF232846.1 | 0 | 100% | 97.12 |
| *Merluccius merluccius* | AF232845.1 | 0 | 100% | 97.12 |
| *Merluccius capensis* | **EF362862.1** | 0 | 100% | 97.12 |
| *Merluccius merluccius* | MT410897.1 | 0 | 100% | 96.88 |
| *Merluccius merluccius* | **EF362874.1** | 0 | 100% | 96.88 |
| *Merluccius merluccius* | **EF362871.1** | 0 | 100% | 96.88 |
| *Merluccius merluccius* | AF232848.1 | 0 | 100% | 96.88 |
| *Merluccius merluccius* | AF232847.1 | 0 | 100% | 96.88 |
| *Merluccius merluccius* | AF232844.1 | 0 | 100% | 96.88 |
| *Merluccius merluccius* | AF232843.1 | 0 | 100% | 96.88 |
| *Merluccius merluccius* | AF232841.1 | 0 | 100% | 96.88 |
| *Merluccius merluccius* | AF232840.1 | 0 | 100% | 96.88 |
| ----- [no other *M. polli* in best 100 blast hits] ----- |  |  |  |  |
| *Merluccius polli* | AY850718.1 | 8.00E-93 | 44% | 99.46 |
| *Merluccius polli* | AY850719.1 | 1.00E-91 | 44% | 98.92 |
| *Merluccius polli* | AY850722.1 | 2.00E-88 | 44% | 98.39 |
| *Merluccius polli* | AF112254.1 | 2.00E-69 | 33% | 100 |
| *Merluccius polli* | AY850721.1 | 4.00E-65 | 32% | 98.55 |
| *Merluccius polli* | AY850720.1 | 4.00E-65 | 32% | 98.55 |

**Figure 2.**
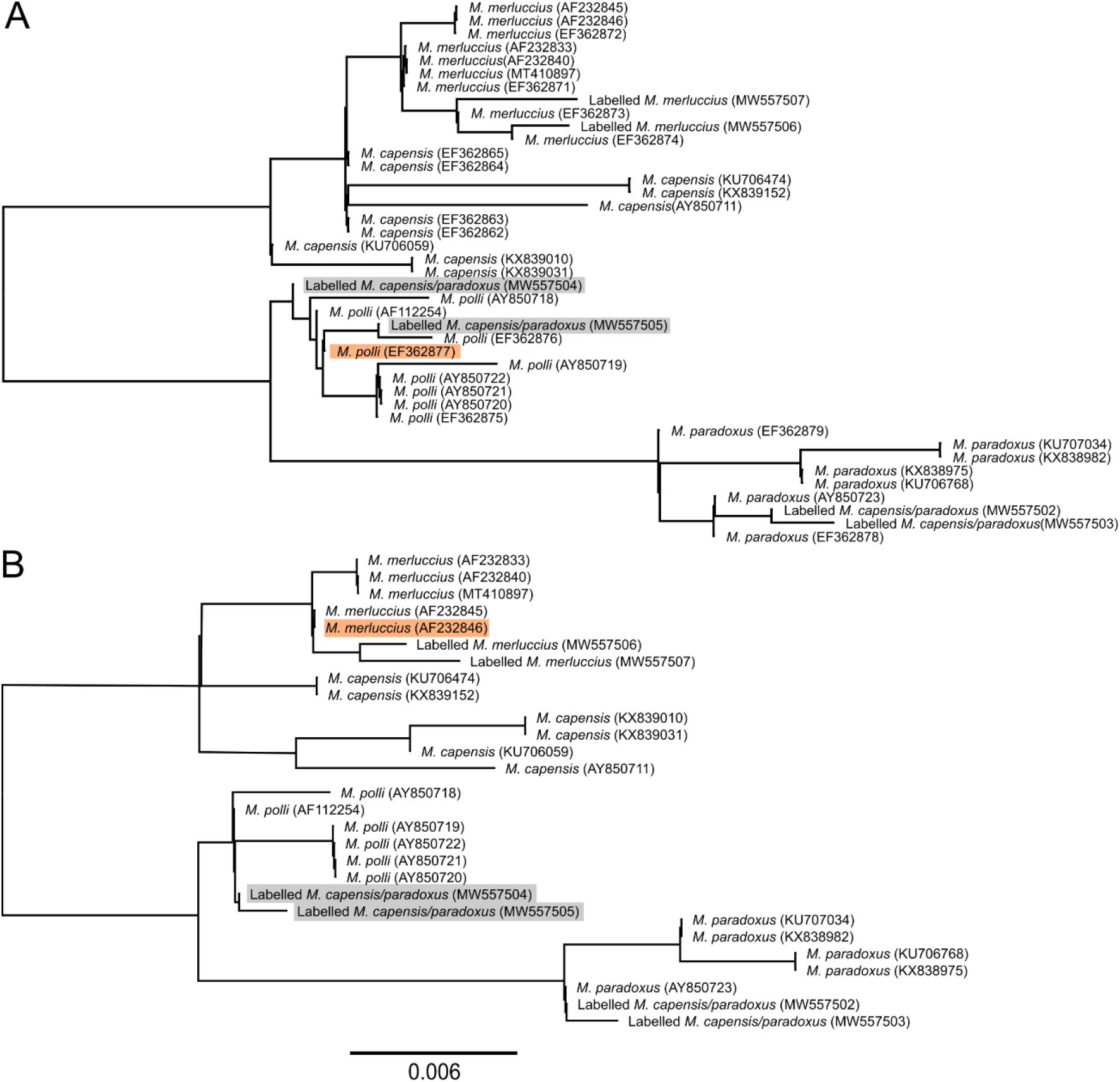
Maximum likelihood phylogenetic inference based on the mitochondrial control region (positions with less than 50% missing data between positions 3 and 433 of EF362862) including all the available *Merluccius* identified sequences from Blanco-Fernandez, et al. ^1^(MW557502-7) and a representative subset of *M. polli*, *M. capensis*, *M. merluccius* and *M. paradoxus* sequences available in GenBank, both including (A) and excluding (B) those from Machado-Schiaffino, et al. ^13^. Sequences to be authenticated and best overall BLAST hit in each case are shaded in grey and orange respectively. The tree is unrooted.

In addition to the above-highlighted potential problems derived from the use of BLAST for species assignment in seafood fraud studies, there is also the issue of contaminated sequences present in GenBank ^14^. As an example, Blanco-Fernandez *et al*.’s work increase the contaminations in GenBank as they have contributed sequences whose taxonomic assignment has relied on best BLAST hit. As consequence of this, there are now three sequences of *T. alalunga* in GenBank labelled *T. thynnus*. These sequences, as well as those not obtained from morphologically identifiable specimens, should be retracted from GenBank to avoid further ramifications.

## Outlook

In summary, we conclude that best BLAST hit cannot be used as evidence of fraud, and that studies on seafood authentication should consider the evolutionary context of the species under study. Not doing so can derive in serious consequences as illustrated by the problems we found in the work of Blanco-Fernandez *et al*., who base their strong claims on tuna mislabeling tends in Spain on erroneous taxonomic assignments.

## Methods

Nucleotide sequences were downloaded from GenBank using provided accession numbers and aligned using ClustalW ^15^. Positions with more than 50% of missing data were removed using Bioedit ^16^ and used for building maximum likelihood phylogenetic trees using DNAML ^17^. BLAST ^18^ searches were performed against the NCBI nucleotide collection (nr/nt) using default parameters.

## Data availability

All data analysed for this reply have been downloaded from GenBank using the accession numbers provided in the Figures.

## Author contribution statement

NDA and NRE conceived the idea, conducted the analyses, and wrote and edited the manuscript.

